# GAGAM: a genomic annotation-based enrichment of scATAC-seq data for Gene Activity Matrix

**DOI:** 10.1101/2022.01.24.477458

**Authors:** Lorenzo Martini, Roberta Bardini, Alessandro Savino, Stefano Di Carlo

## Abstract

Single-cell Assay for Transposase Accessible Chromatin using sequencing (scATAC-seq) is rapidly becoming a powerful technology to assess the epigenetic landscape of thousands of cells. However, the current great sparsity of the resulting data poses significant challenges to their interpretability and informativeness. Different computational methods are available, proposing ways to generate significant features from accessibility data and process them to obtain meaningful results. In particular, the most common way to interpret the raw scATAC-seq data is through peak-calling, generating the peaks as features. Nevertheless, this method is dataset-dependent because the peaks are related to the given dataset and can not be directly compared between different experiments. For this reason, this study wants to improve on the concept of the Gene Activity Matrix (GAM), which links the accessibility data to the genes, by proposing a Genomic-Annotated Gene Activity Matrix (GAGAM), which aims to label the peaks and link them to the genes through functional annotation of the whole genome. Using genes as features solves the problem of the feature dataset dependency allowing for the link of gene accessibility and expression. The latter is crucial for gene regulation understanding and fundamental for the increasing impact of multi-omics data. Results confirm that our method performs better than the previous GAMs.

## 1 Introduction

Recent advances in New Generation Sequencing (NGS) technologies paved the way for single-cell multi-omics data analysis, which captures different facets of cells’ regulative state, including the epigenome, the genome, the transcriptome, and the proteome [19]. Multi-omics approaches increase resolution and sensitivity in the characterization of cellular states, the identification of known or new cellular phenotypes, and the understanding of cell dynamics [13]. This characteristic supports a quantitative and comprehensive approach to study cellular heterogeneity [5].

In particular, the combination of transcriptomic and epigenomic data provides integrated information on the functional activation of genes and the structural organization of chromatin. There are different experimental approaches to generate epigenomic data. These includes accessibility measurements, which indicate whether chromatin is open or closed at genomic locations, exposing other genomic regions for transcriptional and regulatory processes [15]. These data have a very different organization than transcriptomic data indicating the expression level of genes.

Analyzing data from multiple omics does not directly imply to gain richer information on the cellular system, nor to gain a systemic understanding of regulative modalities generating the data. To achieve that, a multi-omics analysis must combine data-driven and model-driven approaches by considering not only the multiple modalities but also their interrelations in the cellular system [29]. To consider them together, it is necessary to correlate the expression level of genes (i.e., transcriptomic analysis) and the accessibility of their relevant coding and regulatory genomic regions.

The concept of gene activity, i.e., the overall accessibility of a gene allowing its transcription inside the cell [26], facilitates comparison between accessibility and expression data. Gene activity is a necessary but not sufficient condition to transcript a gene: a cell can have a coding region accessible at the epigenomic level and the corresponding gene either strongly, weakly, or not expressed at all at the transcriptomic level. This must be considered when comparing transcriptomic and epigenomic data and build approaches to analyze them jointly.

A Gene Activity Matrix (GAM) [26] is an effective way to summarize accessibility information deriving from single-cell experiments. In a GAM, columns identify cells while rows identify genes. An element of the matrix (*GAM_g,c_*) represents the Gene Activity Scores (GAS) of the gene *g* in cell *c* [26]. The GAS is a value describing the activity of a gene in a cell in a given model. The use of the same genes in expression and activity experiments makes transcriptomic data directly comparable with epigenomic data.

Current approaches to compute GAMs derive primarily from data-driven strategies, which show limitations in capturing the contextual meaning and the regulative implications of epigenomic data. This work takes a step towards integrating transcriptomic and epigenomic data to support consistency in the joint consideration of gene activity and gene expression. In particular, this paper introduces a data- and model-driven computation of a Genomic Annotated GAM (GAGAM), which leverages accessibility data and information from genomic annotations of regulatory regions to weigh the gene activity with the annotated functional significance of accessible regulatory elements linked to the genes. GAGAM helps improve the resolution, explainability, and interpretability of the results of the clustering and differential activity analyses, supporting the study of cellular heterogeneity based on epigenomic data alone [26].

## 2 Background

Single-cell Assay for Transposase Accessibility Chromatin sequencing (scATAC-seq) is rapidly becoming the primary way to assess the accessibility of the whole genome at the single-cell resolution. ScATAC-seq datasets employ different ways to define meaningful features to allow their analysis, as shown in [9]. One of the most popular is the “peak calling”, which defines peaks (i.e., intervals on the genome that have a local enrichment of transposase cut-sites) from an experiment-dependent set of chromosomal regions [36]. Since resulting peaks directly derive from the experimental results, they are not univocal, as in transcriptomic data. This hampers comparison of different analyses results and identification of cell-type related marker genes.

As described before, a GAM is an effective way to define robust accessibility features. The GAM considers the overall accessibility of the genomic regions linked to a gene. Using scATAC-seq data, the gene activity scores composing the elements of a GAM can be computed as the accessibility of the peaks related to a gene in a cell. However, the way to link peaks to the correct genetic region on the genome is not unique, and in the literature, there are three main strategies:

1. The GeneScoring sums the peaks in a broad region before and after a gene’s Transcription Starting Site (TSS), weighted by their distance from it [18]. This is the easiest way to define the activity of a gene, but it does not consider all the regulatory aspects.
2. Cicero defines the activity of a gene as the accessibility of the peaks overlapping the TSS and the accessibility of all the co-accessible peaks [26]. This method is more structured than the previous one. However, it identifies the genes through a single DNA base, i.e., the TSS, limiting the effectiveness of the approach. Moreover, co-accessibility evaluation is a very long and computationally heavy process, and the GAS estimation does not consider the meaningfulness of the peak.
3. Signac GAM counts all the raw reads in the gene body [28]. The main limitation of this method is the necessity of a fragment file related to the dataset, which contains all the fragments read in each cell. It is a large file and rarely available, thus making the computation often impossible.

In general, all these methods oversimplify the relationship between a gene and its accessibility. The epigenetic mechanisms are related to the regulation and resulting expression of the genes. However, this association is not direct and linear. If a gene is accessible, it is not necessarily also expressed: the association only gives an insight on whether transcription is possible or not.

Studying how accessibility links to gene expression becomes relevant due to the emergence of new multi-omic Next-Generation Sequencing (NGS) techniques allowing performing both scATAC-seq and Single-cell RNA sequencing (scRNA-seq) simultaneously. One way to achieve multi-omic consistent integration is to employ a model-driven approach.

For this reason, GAGAM introduces a new way to construct a GAM based on the functional annotation of the peaks. GAGAM is not only a new GAM but also a new way to interpret epigenomic data in perspective to link them with transcriptomic data. This method employs publicly available genomic annotations and evaluates the activity based on the regulatory elements linked to the gene by elaborating only the peaks related to the genes and their regulatory regions. Thus, it provides a model-driven GAS that reflects the accessibility to the whole transcription machinery, drawing a direct link to the gene expression.

## 3 Materials and Methods

Fig. 1 introduces the workflow for computation and evaluation of GAGAM starting from a scATAC-seq dataset.

**Fig. 1.**
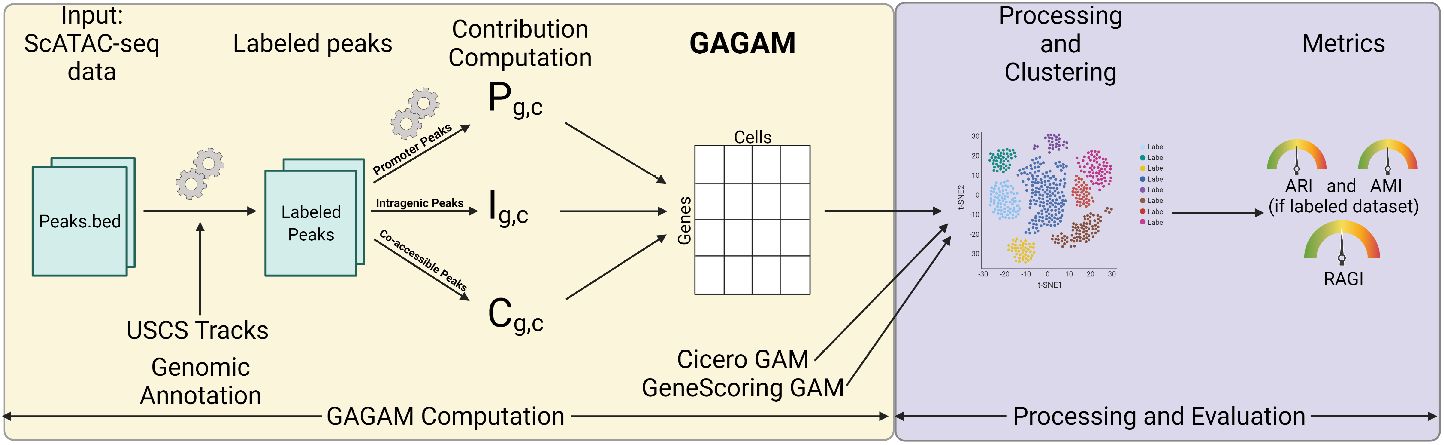
Workflow for computation and evaluation of GAGAM. The workflow starts with scATAC-seq data, and labels the peaks with the help of genomic annotations and USCS tracks. Then it computes the three contributions forming GAGAM. GAGAM is evaluated and compared to other GAMs through clustering experiments with three well-established metrics: Adjust Rand Index (ARI) [17], and Adjust Mutual Information (AMI) [35] (if the dataset is labeled) or Residual Average Gini Index (RAGI) [6] (if the dataset is not labeled).

A scATAC-seq dataset contains a set *P* of peaks observed in a group of *C* cells. Each peak corresponds to a region of the target genome and is defined by its chromosome and a genomic coordinate pair *p* = (*ch, start, stop*). The dataset is a binary matrix **D**_|*P*|×|*C*|_ where rows are associated with peaks and columns with cells. An element of **D** equal to 1 denotes a peak (row) accessible in a cell (column).

The main contribution of GAGAM is to exploit information regarding overlaps of peaks, gene bodies, and genetic regulatory regions (i.e., promoters and enhancers) to build a GAM with higher information content.

### 3.1 Genomic Annotation

The genomic annotation of peaks is the first step to constructing GAGAM. This process aims to enrich information regarding peaks with data coming from different genomic annotations useful for a model-driven construction of a GAM. Fig. 2 shows the genomic model considered in this work. It represents a genomic unit, which includes three parts: (i) the gene body region, starting at the Transcription Starting Site (TSS), (ii) the gene Promoter, preceding the coding region, and (iii) a set of Enhancers that are distal to the gene.

**Fig. 2.**
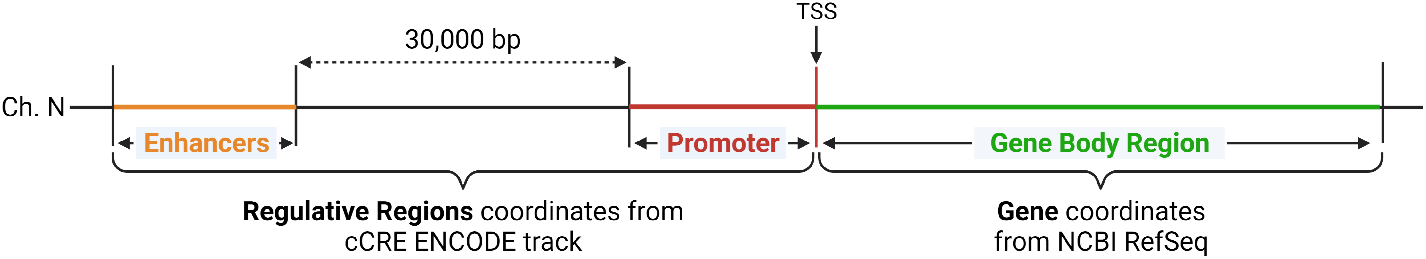
Genomic Model. The genomic model consists of the coordinates of all the genomic regions related to the gene. The gene body region (in green) comes from the NCBI RefSeq Genes annotations. The regulative regions, i.e., Promoter (in red), and Enhancers (in orange), come from the cCRE ENCODE tracks.

The gene coding region is defined using NCBI RefSeq Genes [25] annotations, consisting of genes’ genomic coordinates. Therefore, a gene *g* in a target genome *G* is a tuple defining the gene’s chromosome and its genomic coordinates pair (i.e., *g* = (*ch, start, stop*)). The NCBI RefSeq annotations are accessible using the NCBI Eukaryotic Genome Annotation Pipeline [30]. It consists of an annotated and curated information list of protein-coding and non-protein-coding genes. The annotation also includes all the pseudogenes and miRNA regions. Since GAGAM aims to obtain something as close as possible to the transcriptomic information, it only considers the protein-coding and lncRNA regions.

The regulative genomic regions are elements on the DNA footprints for the trans-acting proteins involved in transcription, either for the positioning of the basic transcriptional machinery or for the regulation. The annotation tracks are associations between a genomic region and a label indicating the function of the region. Given a target genome *G* it is possible to define a set *R* of regulative regions with each region defined by the corresponding chromosome, the genomic coordinates pair, and a label (e.g., promoter or enhancer) indicating the function of the region (*r* = (*ch, start, stop, l*)).

Information regarding regulative gene regions are available from the Encyclopedia of DNA Elements (ENCODE) project, which provides an extensive collection of cell- and tissue-based repertoires of genomic annotations, including, for example, transcription, chromatin organization, epigenetic landscape dynamics, and protein binding sites from the mouse and human genomes [24]. ENCODE data are available through the ENCODE data portal [11].

This work only considers genomic annotations relative to promoter and enhancer functions. These regulatory elements are derived from the ENCODE candidate cis-Regulatory Elements (cCREs). cCREs provides an extensive collection of annotated regions for the human and mouse genomes. Classification of cCREs is based on biochemical signatures, considering DNase hypersensitivity, histone methylation, acetylation, and CTCF binding data [24]. Since this work aims to label peaks from both human and mouse datasets, cCREs tracks (in BigBed format [16]) were collected from ENCODE for both the human [33], and mouse [34] genomes.

The goal of the genomic annotation process is to associate each peak *p* ∈ *P* obtained from a scATAC-seq dataset **D** to a set of genomic annotation labels by analyzing how the peak overlaps to the different genomic regions.

GAGAM labels each peak *p* ∈ *P* with four possible labels: (1) `prom` for peaks overlapping a promoter region, (2) `enhD` for peaks overlapping a distal enhancer region and not a promoter region, (3) `intra` for the peaks contained into a gene body region, and (4) `empty` in all other cases. The rule to assign the label is summarized in the following equation:

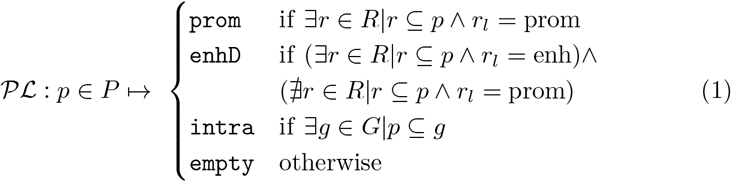

The operator *a* ⊆ *b* is used here to denote that the two regions *a* and *b* belong to the same chromosome with *b* overlapping *a* (i.e., *a_start_* ≥ *b_start_* ∧*a_end_* ≤ *b_end_*). The computation of the intersection between peaks and annotation regions leverages the `bigBedToBed` tool from ENCODE [32].

Performing genome annotation for the mouse genome is straightforward for all considered datasets since both datasets, and annotation tracks refer to the mm10 genomic assembly. Differently, for the human genome, the cCREs annotation track is only available for the hg38 genomic assembly, while the human dataset is based on the hg37 genomic assembly. For this reason, this work leverages the UCSC `LiftOver` tool [20] to convert the peaks’ coordinate ranges from the hg37 to the hg38 assemblies before performing peak labeling.

Given the list of annotated peaks, GAGAM builds a gene activity matrix as a weighted sum of three separated matrices: (i) the promoter peaks matrix (**P**) indicating accessibility of genes associated with promoter peaks, (ii) the intragenic peaks matrix (**I**) indicating the accessibility of genes containing intragenic peaks, and (iii) the co-accessibility matrix (**C**) indicating the accessibility of genes associated with distal enhancer peaks, obtaining a final curated and model-driven evaluation of the activity of the genes:

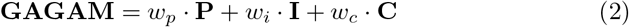

### 3.2 Promoter Peaks Matrix

The promoter peaks matrix exploits model-driven information about promoter peaks to identify relevant genes in the GAM. The golden rule applied in GAGAM is that *a gene in a cell is active if, and only if, its promoter peak is accessible*. This rule reduces the set of interesting genes to consider when constructing a GAM.

To follow this rule, let us denote with *P^p^* ⊆ *P* the subset of peaks in the dataset **D** annotated as promoters (i.e., 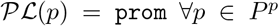) and with 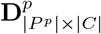 the submatrix of **D** including only rows associated to promoter peaks.

GAGAM constructs a binary matrix **GP**_|*G^p^*|×|*P^p^*|_ associating the set of genes with active promoter peaks (*G^p^*) to their related peaks. To associate a promoter peak to a gene, GAGAM considers the overlapping of an enlarged gene body region including 500bp before the TSS (i.e., an approximation of the mean peak length) with the peak region. Based on this, the promoter peaks matrix is a binary matrix computed as:

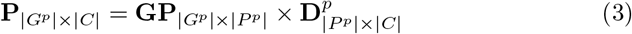

This matrix is a GAM including accessibility data for the subset of genes associated with the promoter peaks. In this way, GAGAM leverages available knowledge on transcriptional regulatory regions to define the active genes based on a model taking into account the knowledge of gene regulation and transcription.

### 3.3 Intragenic Peaks Matrix

GAGAM also considers the contribution of the intragenic peaks (i.e., peaks located in the gene body region) to the overall gene activity score. Similarly to what described before, let us denote with *P^i^* ⊆ *P* the subset of peaks in the dataset **D** annotated as intragenic (i.e., 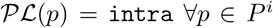) and with 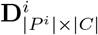 the submatirx of **D** including only rows associated to intragenic peaks.

GAGAM constructs a matrix **GI**_|*G^p^*|×|*p^i^*|_ associating genes with active promoter peaks (*G^p^*) to their related intragenic peaks. This matrix only considers genes with active promoter peaks to follow the GAGAM golden rule (section 3.2). Some of the identified intragenic peaks could be part of genes that do not have a promoter peak. Moreover, it could happen that given a gene region inside a cell, intragenic peaks could be accessible even if the promoter peak is not.

Statistically, there will be more peaks inside the gene body region of a long gene, meaning it might have a higher score after its length. To prevent this bias, GAGAM employs a strategy from the GeneScoring [18] method to compute the elements of **GI**. It weighs the contribution of the intergenic peaks with an exponentially decaying function of their distance from the TSS (i.e., 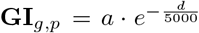 where *a* = 1 if *p* ⊆ *g*, 0 otherwise and *d* is the distance of the peak from TSS). In this way, very long genes are not over-represented because the most crucial part of the gene’s activity is near the promoter. Therefore, the peaks near it are weighted more. Based on this, the intragenic peaks matrix is a matrix computed as:

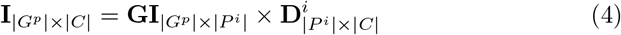

### 3.4 Promoter-Enhancer co-accessibility Matrix

The co-accessibility matrix accounts for the connections between promoters and enhancers. It leverages Cicero [26] to calculate the co-accessibility of the peaks. The co-accessibility represents how couples of peaks tend to be simultaneously accessible in the cells, expressing it in a range between 0 and 1. This calculation “*connects regulatory elements to their putative target genes*” [26], meaning it can find connections between the promoters and the distal regulatory regions where different elements like Transcriptional Factors (TF) bind and enable the transcription.

The first step to compute this matrix is to calculate the co-accessibility from the scATAC-seq data with the Cicero function `run_cicero` (for the explanation of the calculation, refer to [26]). The result is a list of peaks couples with their co-accessibility value (*ca*) and distance (d) in the form *conn* = (*p*^1^, *p*^2^, *ca,d*).

GAGAM selects only couples of promoter-enhancer peaks, i.e., couples with *p*^1^ ∈*P^p^* and *p*^2^ ∈ *P^e^* (or vice versa), with *P^e^* ⊆ *P* representing the subset of peaks in the dataset **D** annotated as enhancers (i.e., 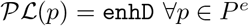).

Moreover, GAGAM keeps only couples with *ca* ≥ *ca_m_* (with *ca_m_* the mean value of all the co-accessibility scores above zero) and *d* ≤ *d_th_* (with *d_th_* = 30, 000 *bp* the distance threshold defined as suggested by the guidelines of Cicero [26]).

To calculate the co-accessibility matrix **C**, GAGAM uses three matrices. First, the binary matrix **GP**_|*G^p^*|×|*p^p^*|_ previously defined in section 3.2 and associating genes with promoter peaks. Second, the matrix **PE**_*P^p^*|×|*p^e^*|_ associating promoter peaks and enhancer peaks. The elements of this matrix are the coaccessibility values *ca* of the couples of peaks available in the list produced by Cicero, and 0 otherwise. Third, the matrix 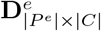 is a submatrix of **D** including only rows associated with enhancer peaks.

Based on this, the co-accessibility matrix is computed as:

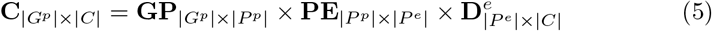

## 4 Results and discussion

### 4.1 Evaluation strategy

This section evaluates GAGAM by looking at different aspects.

First, GAGAM represents an interpretation of scATAC-seq data. As the majority of single-cell experiments, it must identify cellular heterogeneity. Based on this consideration, the first approach to evaluate the capabilities of this new gene activity matrix is to employ one of the many available pipelines to process GAMs (Fig. 1). This work uses Monocle3 [31], given its simplicity and the fact that Cicero GAM (see section 2) is dependent on it.

The standard Monocle workflow starts with a GAM, performs Principal Component Analysis (PCA), visualizes the cells in 2D using UMAP [23], and most importantly, performs cells clustering. The clustering results should at least partially represent the cellular heterogeneity of the dataset. This can be measured using a group of metrics thoroughly discussed in section 4.2.

Moreover, it can be proved that GAGAM and, in particular, the selected genes are not just a product of data manipulation but are biologically meaningful in two ways. First, performing differential activity analysis on the GAM can show that the differentially active genes are cell-type specific. This would also demonstrate that employing marker genes allows classifying scATAC datasets, something not possible with raw data. Second, using the RAGI index (one of the metrics for the evaluation of the clustering performances defined in section 4.2), it is possible to assess the informativity of the GAGAM.

### 4.2 Metrics definition

The evaluation strategy proposed in section 4.1 is based on unsupervised clustering of cells based on the selected genes. The obtained clusters are the outputs that must be analyzed to understand if they represent cell heterogeneity. There are two scenarios: (i) the starting dataset has cell labels; thus, each cell has a label identifying its cell type, or (ii) there are no available cell labels, so there is no ground truth to compare.

In the first case, the most direct way to measure the quality of the clustering process is to compare the clusters to the cell-type labels. To show how much the two classifications are similar, this paper uses the Adjust Rand Index (ARI) [17] and Adjust Mutual Information (AMI) [35] from information theory. These two metrics are often employed for this type of evaluation. In particular, [6] uses them for their benchmarking. Thus, they help compare their results with those produced in this paper. ARI and AMI range between 0 and 1, where 1 is a perfect match, and 0 is complete uncorrelation. This evaluation employs the R package ARICODE [8], which easily allows their calculation.

The second case requires a different approach. Since there is no reference classification, ARI and AMI cannot be used. One method is to calculate ARI and AMI comparing the results with the clustering-based labels obtained from the scATAC-data data processing. Otherwise, [6] proposes a very fitting way: the Residual Average Gini Index (RAGI) [6]. The RAGI investigates the differences in the Gini index of markers and housekeeping genes. The idea is that a good clustering should have marker genes active only in specific clusters and housekeeping genes over all the cells. Therefore, RAGI can measure the quality of the GAM itself. A good GAM should convey meaningful biological information that should translate into a difference between the two sets of genes. Therefore, the RAGI estimates if a GAM can correctly assess the gene activity. Anyway, RAGI has the problem of being highly dependent on the employed genes to calculate it. Still, the concept of housekeeping genes and, even more, marker genes are not well-defined [22]. Therefore, it is essential to carefully choose the right set of marker genes strictly related to the dataset sample. In this work, the list of housekeeping genes derives from [10] for humans and [12] for the mouse. On the other hand, the marker genes list comes from the CellMarker [37] database, which provides a curated list of markers per tissue. For the mouse brain datasets, this work also employs markers from [14], and [21].

### 4.3 Datasets

GAGAM was tested on five datasets (see Table 1). Two datasets are from the 10XGenomic platform [27], and consist of a collection of respectively 5,335 (*10X V1.0.1 PBMC* [1]) and 4,623 (*10X V2.0.0 PBMC* [3]) cells from human Peripheral Blood Mononuclear Cells (PBMC) samples. From the 10XGenomic platform, there is also a mouse brain dataset with 5,337 cells (*10X V1.1.0 Brain* [2]). All three datasets do not have cell labels. Therefore, ARI and AMI evaluations are applicable only on the clustering-based labels. Next, this study employed a dataset of bone marrow (*Buenrostro2018*) from [4]. This dataset consists of 2,034 cells and provides cell-type classification. The last dataset comes from a multi-omic SNARE experiment (*SNARE* [7]). It consists of 10,309 cells from the mouse cortex and comes with a partial classification of the cells. Two of the considered datasets (*10X V1.0.1 PBMC* and *Buenrostro2018*) derive from [6], a paper performing a benchmarking analysis on different methods allowing for easy comparison of results.

**Table 1.** Datasets employed.

| Dataset | Species | Tissue | Cells labels | Reference |
| --- | --- | --- | --- | --- |
| *10X V1.0.1 PBMC* | Human | PBMC | No | [1] |
| *10X V2.0.0 PBMC* | Human | PBMC | No | [3] |
| *10X V1.1.0 Brain* | Mouse | Brain Cortex | No | [2] |
| *Buenrostro2018* | Human | Bone Marrow | Yes | [4] |
| *SNARE* | Mouse | Brain Cortex | Yes | [7] |

### 4.4 Results

This section compares the performance of GAGAM with two state-of-the-art GAM computation pipelines (i.e., Cicero and GeneScoring) following the evaluation strategy proposed in section 4.1. Since GAGAM is constructed from three contributions (see eq. 2), it is advisable to evaluate different combinations to select the best one. This experimental setup considers two versions of GAGAM: GAGAM1 constructed considering only the promoter peaks and the co-accessibility (i.e., *w_p_* = 1, *w_i_* = 0, and *w_c_* = 1) and GAGAM2 created using the complete GAGAM workflow (i.e., *w_p_* = 1, *w_i_* = 1, and *w_c_* = 1).

Figures 3 and 4 reports AMI and ARI results comparing *Buenrostro2018* [4] and *SNARE* [7] clusters, with their ground truth labels, and the other datasets against the scATAC clustering results. Overall, Figures 3 and 4 shows that both versions of GAGAM perform equally or better than Cicero and GeneScoring. Only on *10X V1.1.0 Brain*, GeneScoring has a higher metric value than GAGAM. However, comparing the results in [4] with the ones reported in [6] on the same dataset shows how the GAGAM performances are on the high end of the bench-marked paper methods (as shown in Table 5 from [6]).

**Fig. 3.**
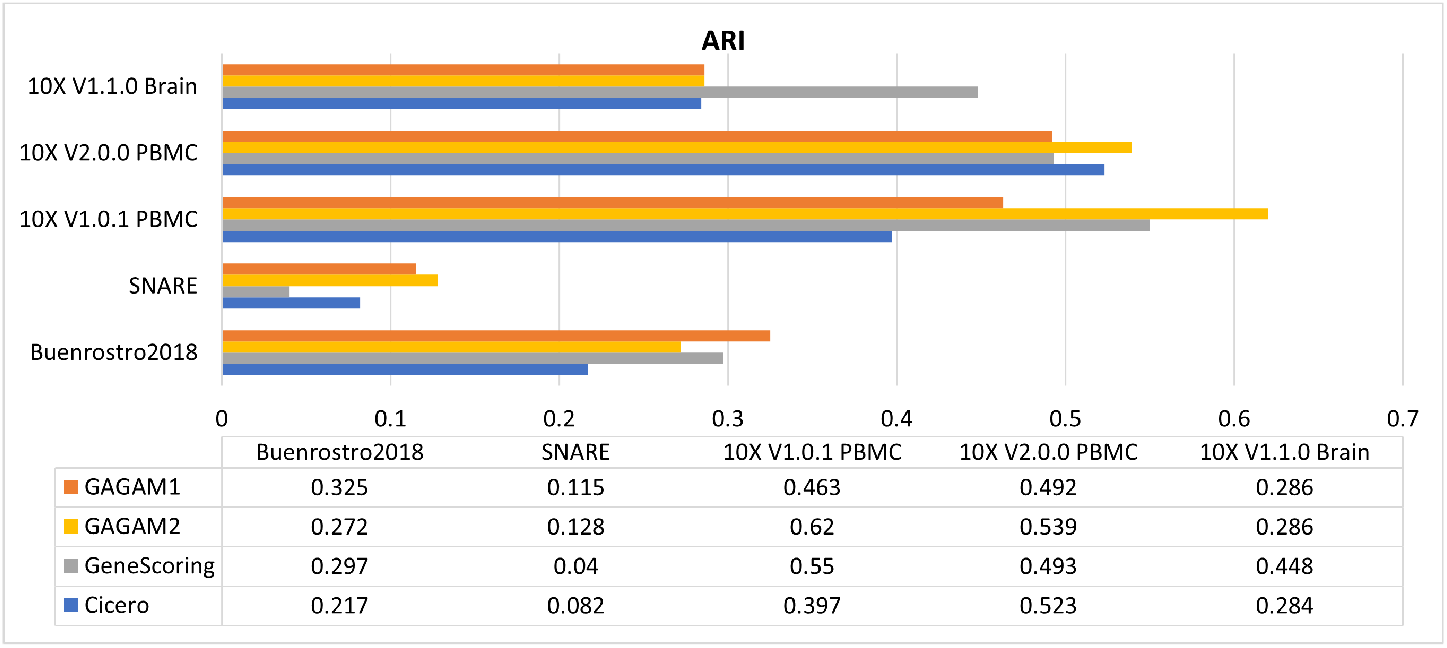
ARI results of the four methods for the five datasets.

**Fig. 4.**
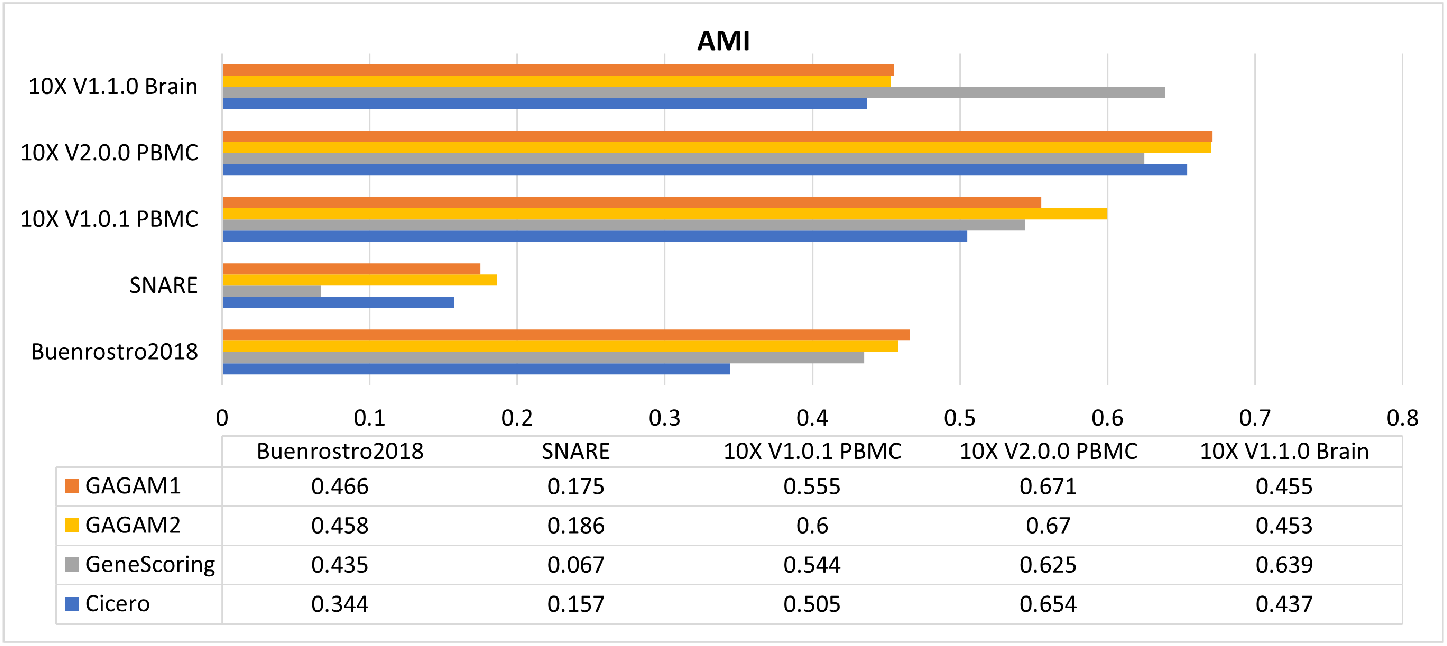
AMI results of the four methods for the five datasets.

Next, RAGI has been computed for all datasets to assess the clustering results and the information content of the GAMs. The results are in Figures 5 and 6. For the three different types of tissues, we employed three different sets of curated markers, while the housekeeping genes were shared between the same species datasets. For each method and dataset, there are two different results. One is the RAGI score calculated on each GAM concerning the clustering results. The other is computed on each GAM but resorting to the cell labels (when available) or the clustering-based labels obtained from the scATAC data processing. This way, all methods are evaluated against the same partition to understand which GAM is the most biologically consistent. In particular, the *10X V1.0.1 PBMC* dataset is assessed with this metric in [6], and GAGAM outperforms all the methods illustrated there.

**Fig. 5.**
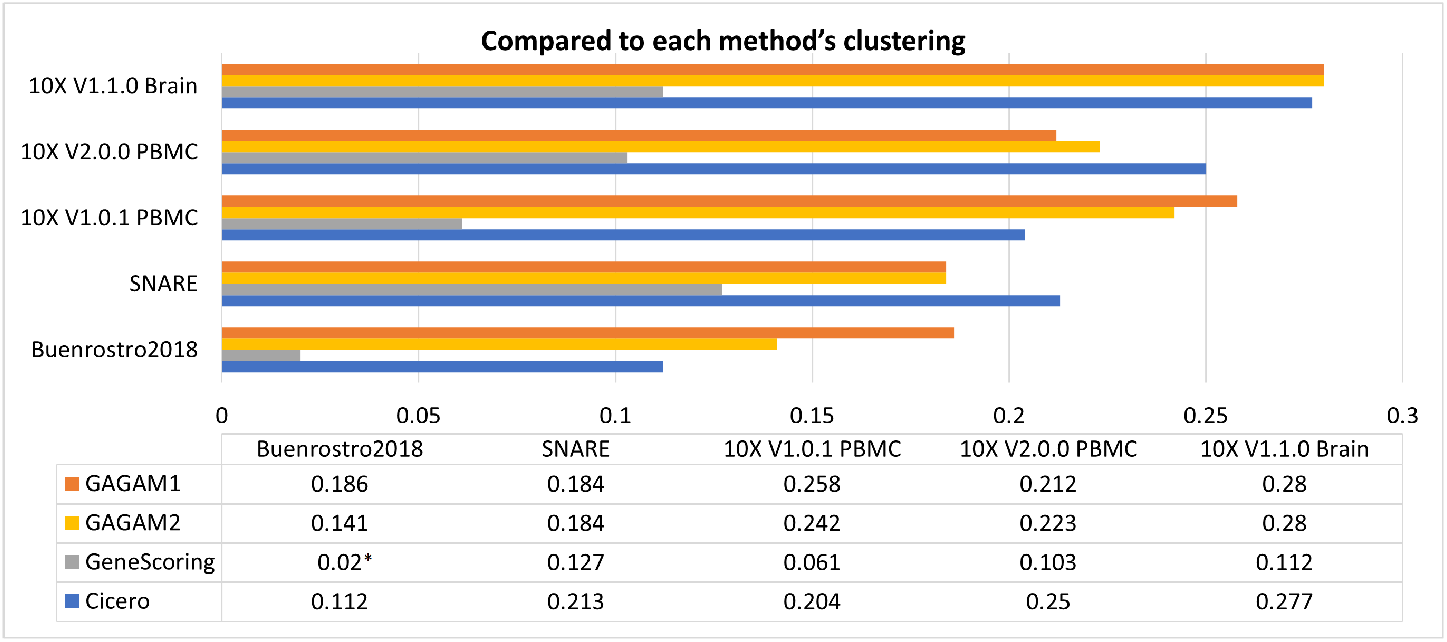
RAGI results of the four methods for the five datasets, when compared to each method’s clustering.

**Fig. 6.**
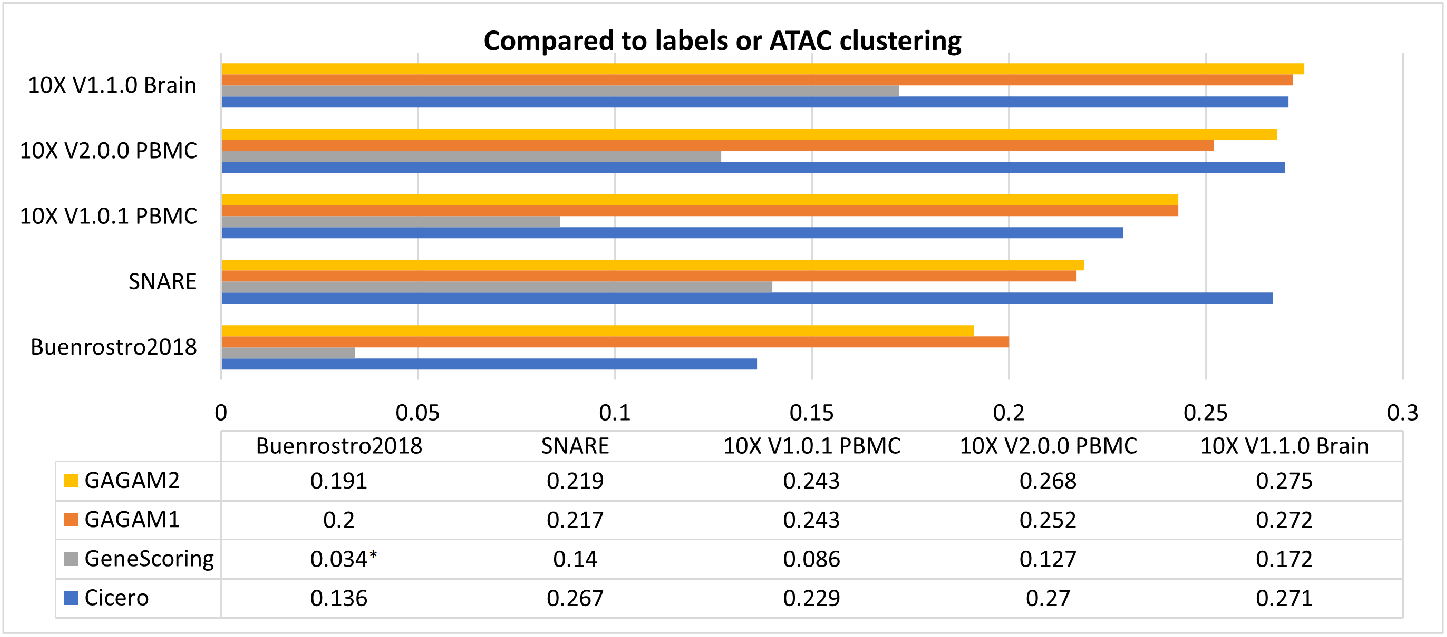
RAGI results of the four methods for the five datasets, when compared to labels or ATAC clustering.

In general, the results show how GAGAM has consistently good performances. Nevertheless, in this case, Cicero performs better on the *SNARE* dataset. Instead, GeneScoring offers low performances. Although its clustering results are consistent with the ground-truth classification (as indicated by ARI and AMI), the actual scores are not well defined. This suggests the importance of evaluating the GAMs on both metrics. Therefore, although there are some cases where Cicero and GeneScoring have better results than GAGAM, the latter has a consistent behavior on all the metrics, meaning it is the most reliable method on both clustering results and actual GAS computation. It is essential to highlight that some of the RAGI results (marked with *) have a p-value over the tolerable threshold (0.05), so they are not statistically meaningful, but we report them anyways.

## 5 Conclusions

In conclusion, GAGAM is a new method to obtain a Gene Activity Matrix from scATAC-seq data. It is based on a model-driven approach leveraging genomic annotations of genes and functional elements. It introduces the promoter peak accessibility into the score, which is necessary for the gene’s activity. Then, it considers the contribution of intragenic peaks, weighted by their distance from the TSS and the enhancer peaks connected to the promoter. The score obtained this way represents a good model of the gene activity interpreted as the set of elements that should be accessible to allow gene transcription.

Experimental results demonstrate how GAGAM generally performs better against other GAMs concerning its ability to identify cellular heterogeneity. Specifically, the clustering obtained from GAGAM is evaluated with ARI, AMI, and RAGI and has better results than Cicero and GeneScoring on all of these metrics. In addition, GAGAM is a suitable method to interpret accessibility data in general. Indeed, since it employs genes as features, it allows analyzing scATAC-seq data through well-studied and investigated concepts like marker genes. The same analysis would not be possible with raw accessibility data. RAGI results support this claim and highlight the activity differences between marker and housekeeping genes. This activity proves that the features selected in GAGAM (i.e., the genes) and their activity scores are biologically meaningful. Therefore, GAGAM provides an optimal and reliable middle ground between the accessibility data and the gene expression data, crucial for future works in a field where multi-omics single-cell techniques are fastly growing.

In conclusion, GAGAM is a promising and reliable way to interpret scATAC-seq data, which focuses on the accessibility of the genes and their regulatory elements, acting as a direct link between epigenomic and transcriptomic.

